# Disruption of metapopulation structure effectively reduces Tasmanian devil facial tumour disease spread at the expense of abundance and genetic diversity

**DOI:** 10.1101/2021.10.04.463046

**Authors:** Rowan Durrant, Rodrigo Hamede, Konstans Wells, Miguel Lurgi

**Author notes:** **Corresponding author:** Miguel Lurgi. Room 225BA. Wallace Building. Department of Biosciences, Swansea University. Singleton Park, Swansea SA2 8PP, UK. Institute of Biodiversity, Animal Health and Comparative Medicine, College of Medical, Veterinary and Life Sciences, University of Glasgow, Glasgow G12 8QQ, UK. **Statement of co-authorship:** ML & KW conceived the study. All authors designed the research. RH contributed empirical data to guide modelling. RD performed modelling work and analysed output data. RD wrote the first draft of the manuscript, and all authors contributed substantially to revisions. **Data accessibility statement:** All the source code developed to implement the model and run the model simulations to produce the results presented in this study has been uploaded to the GitHub repository: https://github.com/RowanDurrant/DFTD.

## Abstract

Metapopulation structure (i.e. the spatial arrangement of local populations and corridors between them) plays a fundamental role in the persistence of wildlife populations, but can also drive the spread of infectious diseases. While the disruption of metapopulation connectivity can reduce disease spread, it can also impair host resilience by disrupting gene flow and colonisation dynamics. Thus, a pressing challenge for many wildlife populations is to elucidate whether the benefits of disease management methods that reduce metapopulation connectivity outweigh the associated risks.

Directly transmissible cancers are clonal malignant cell lines capable to spread through host populations without immune recognition, when susceptible and infected hosts become in close contact. Using an individual-based metapopulation model we investigate the effects of the interplay between host dispersal, disease transmission rate and inter-individual contact distance for transmission (determining within-population mixing) on the spread and persistence of a transmissible cancer, Tasmanian devil facial tumour disease (DFTD), from local to regional scales. Further, we explore population isolation scenarios to devise management strategies to mitigate disease spread.

Disease spread, and the ensuing population declines, are synergistically determined by individuals’ dispersal, disease transmission rate and within-population mixing. Low to intermediate transmission rates can be magnified by high dispersal and inter-individual transmission distance. Once disease transmission rate is high, dispersal and inter-individual contact distance do not impact the outcome of the disease transmission dynamics.

Isolation of local populations effectively reduced metapopulation-level disease prevalence but caused severe declines in metapopulation size and genetic diversity. The relative position of managed (i.e. isolated) populations within the metapopulation had a significant effect on disease prevalence, highlighting the importance of considering metapopulation structure when implementing metapopulation-scale disease control measures. Our findings suggests that population isolation is not an ideal management method for preventing disease spread in species inhabiting already fragmented landscapes, where genetic diversity and extinction risk are already a concern, such as the Tasmanian devil.

## Introduction

Infectious diseases are a major threat to the long-term survival of wildlife populations (De Castro & Bolker 2005). Outbreaks of infectious diseases such as chytridiomycosis, infections caused by the Gram-negative bacteria *Pasteurella multocida*, or white-nose syndrome have caused threatening declines in populations of amphibians (Scheele *et al*. 2019), saiga antelope (Fereidouni *et al*. 2019), and bats (Hoyt *et al*. 2020), respectively. Infectious cancers in wildlife are nowadays recognised as a major conservation problem, particularly for endangered species and populations with restricted distribution (McAloose & Newton 2009; Hamede *et al*. 2020b). Genetic bottlenecks, changes in global weather and anthropogenic activities resulting in exposure to toxins, oncogenic pathogens, immunosuppression, stress and urbanisation have been regarded as contributors to the emergence of cancers in wildlife (Giraudeau *et al*. 2018; Pesavento *et al*. 2018; Ujvari *et al*. 2018). As well as threatening biodiversity, wildlife diseases are increasingly being acknowledged as potential risks to human health and domestic livestock (Meslin 1997; Rodriguez-Morales *et al*. 2020), leading to substantial economic loss and compromising human wellbeing (Grenfell & Gulland 1995; Daszak *et al*. 2000).

Management of wildlife diseases is often carried out at the individual or local population level (Wobeser 2002). However, transmission between local populations is a fundamental mechanism of infectious disease spread at intermediate to large spatial scales (May & Anderson 1979; Colizza & Vespignani 2008; North & Godfray 2017), and has been identified as one of the main causes of epidemics in both humans (Wilson 1995) and animals (Fèvre *et al*. 2006), including the recent global spread of COVID-19 (Shi *et al*. 2020; Wells *et al*. 2020; Zhang *et al*. 2020). Increased host movement drives both the regional spread of disease (Viboud *et al*. 2006; Colizza & Vespignani 2008; North & Godfray 2017), and increases the risk of host extinction from disease in theoretical metapopulation systems (Hess 1996; North & Godfray 2017). However, predicting the influence of metapopulation structure on the dynamics of disease-burdened host populations is not a trivial task, particularly for emerging infectious disease with high mortality. While increasing connectivity may facilitate efficient disease spread and disease-induced host population decline, reduced population connectivity may disrupt vital patch colonization dynamics and the maintenance of host genetic diversity (Fulford *et al*. 2002; Hess 1996). In some instances, highly connected host populations can even be less burdened by disease due to increased levels of resistance facilitated by high gene flow (Jousimo *et al*. 2014).

In practice, the success of containing infectious disease spread at landscape scale may depend crucially on the landscape context within which local populations are connected to each other. Landscape-scale disease control strategies aimed at limiting population connectivity and host movement such as fencing, have been used during outbreaks of, for example, African swine fever and chronic wasting disease (Mysterud & Rolandsen 2019). Restricting the movement of individual animals in order to reduce the spread of disease is tempting, but this could potentially come with great ecological costs, such as disrupting migratory patterns and reducing gene flow between populations, as barriers that reduce disease spread have also been shown to shape landscape genetics (Robinson *et al*. 2013).

Devil facial tumour disease (DFTD) is a transmissible cancer that has decimated populations of Tasmanian devils (*Sarcophilus harrisii*) across nearly the entire geographical range of the species (Hawkins *et al*. 2006; Cunningham et al 2021). The disease is a clonal cancerous cell line originated from a mutated Schwan cell in a female devil in north-eastern Tasmania in the 1990’s (Pearse and Swift 2006; Murchison et al 2010). Transmission of DFTD between individuals occurs when infected and susceptible individuals bite each other, a common behaviour in this species (Hamede *et al*. 2013), and is facilitated by the ability of tumours cells to epigenetically downregulate the host’s Major Histocompatibility Complex (Siddle *et al*. 2013). Prevalence of DFTD is considerably high, (up to 80%), even after populations have suffered significant declines, suggesting that transmission is mostly frequency dependent (McCallum *et al*. 2009), which is driven by the increase in contact rates and biting wounds during the mating season (Hamede *et al*. 2009; Hamilton *et al*. 2019). The spread of DFTD across Tasmania is particularly worrying due to the large declines in numbers associated with it, with an average decline of affected populations by 77% (Lazenby *et al*. 2018) and some local populations having declined by as much as 90% (McCallum *et al*. 2007). Landscape-scale spread of the disease occurred relatively fast, with the disease now found on over 90% of the landmass of Tasmania, and the south-west of the island being the only area where the disease has not been detected (Cunningham et al 2021).

Previous work has generated a good understanding of how DFTD spreads between individuals, how it leads to population decline, and how it drives demographic changes within local populations (Jones *et al*. 2008; McCallum *et al*. 2009; Beeton & McCallum 2012; Bruno *et al*. 2017; Wells *et al*. 2019). However, we still know surprisingly little about the processes that drive the spread of DFTD on a regional scale. Counter to the findings of previous single-population models of DFTD that predicted a high probability of host extinction (McCallum *et al*. 2009), a recent metapopulation modelling study by Siska *et al*. (2018) found that the recolonization of extinct local devil populations in a metapopulation may be strong enough to prevent their complete extinction. This suggests that metapopulation structure, and the population dynamics that play out on this network of populations, might play an important role in maintaining devil populations. The survival of devils in Siska *et al*.’s simulations may be at least in part due to the “rescue effect” (Brown & Kodric-Brown 1977; Gotelli 1991), in which the recolonisation of extinct populations is possible via the influx of individuals from remaining populations (Levins 1969). Given its instrumental role in maintaining population viability, metapopulation structure is thus expected to play a fundamental role on disease transmission and recovery (May & Anderson 1979; Hess 1996; Grenfell & Harwood 1997; Colizza & Vespignani 2008). We thus expect metapopulation dynamics to play a fundamental role in DFTD transmission across devil populations in Tasmania.

Here we aim at unveiling the role of local-scale processes, such as within-population mixing and transmission, and their interaction with regional-scale processes of between-population dispersal, in driving the regional spread of DFTD in structured metapopulations. We developed an individual-based metapopulation model of DFTD spread and investigated the effects of altering the magnitude of within-population mixing, devil-to-devil disease transmission and devils’ dispersal rate on metapopulation disease dynamics. We additionally explored the effects of disrupting metapopulation structure on disease dynamics by isolating local populations. This enabled us to assess whether metapopulation fragmentation via isolation methods such as fencing or geographic barriers would be a viable strategy for managing regional disease outbreaks and how it would affect the size and landscape-scale genetic diversity of devil populations.

We hypothesised that dispersal rate would have the largest influence over regional disease spread, as opposed to within-population mixing and disease transmission probability, by allowing for local outbreaks to become regional. We also predicted that isolating populations (i.e. metapopulation fragmentation) would lead to a decrease in disease prevalence and an increase in devil abundance at metapopulation scales relative to simulations where no populations are isolated; this would however, also cause a decrease in genetic diversity. We predict that these effects would be most pronounced when isolating populations according to their connectivity profile within the metapopulation network (e.g. how well connected they are to other populations).

## Methods

We developed an individual-based metapopulation model of DFTD spread in Tasmanian devils on the island of Tasmania, Australia. Metapopulation structure was defined as a network of local populations (nodes) connected via dispersal corridors (links) for devils to move / disperse across populations. Due to the challenge of incomplete landscape scale devil surveillance across Tasmania, geographical location and size of local devil populations in our metapopulation was determined based on known suitable habitat for the species. Local population and disease transmission dynamics were modelled according to our previously developed model (Wells *et al*. 2019), parameterised by long-term detailed monitoring data (Hamede *et al*. 2015; Wells *et al*. 2017).

Using this model, we performed computer simulations with different combinations of parameter values governing dispersal, inter-individual contact distance, and disease transmission rate to investigate the factors driving regional disease spread. Further simulations were performed to assess the effects of isolating local populations (by removing links from selected nodes) as a management strategy on landscape-scale population size, disease prevalence and genetic diversity.

### Metapopulation structure

We defined metapopulation structure as a network of local populations connected according to their proximity to each other. Geographical location and size of local populations was determined using the position and area of patches of sclerophyll forests and coastal heath across Tasmania, where devil density was known to be reasonably high pre-DFTD to assume viable populations according to available devil surveillance data (Guiler 1970b; Department of the Environment 2020). Land cover data was obtained from the digital vegetation map of Tasmania, TASVEG version 2 (TASVEG2) (Department of Primary Industries, Parks, Water and Environment (Tasmania) 2009). Only habitat patches ≥ 5 km^2^ in size were considered. This procedure removed any patches where a local devil population was unlikely to persist due to restricted habitat availability, while maintaining a connected metapopulation network. Predicted devil density of each local population was extracted from recently published density estimates (Cunningham *et al*. 2021). We overlaid the TASVEG2 vegetation map over Cunningham *et al*.’s density estimates map for the year 1986 (10 years prior the estimated appearance of DFTD in Tasmania, and the starting point of our simulations) and extracted the mean devil density over the area of each habitat patch. These local population estimates were used as the initial population numbers at the start of the model simulations. Any populations with a devil density below a threshold of 0.5 devils/km^2^ were excluded from the metapopulation. This yielded a total of 477 populations, with a mean habitat patch area of 13.3 km^2^ (minimum area 5.0 km^2^, maximum area 102.2 km^2^) (Fig. 1; Supp. Fig. S1B).

**Figure 1.**
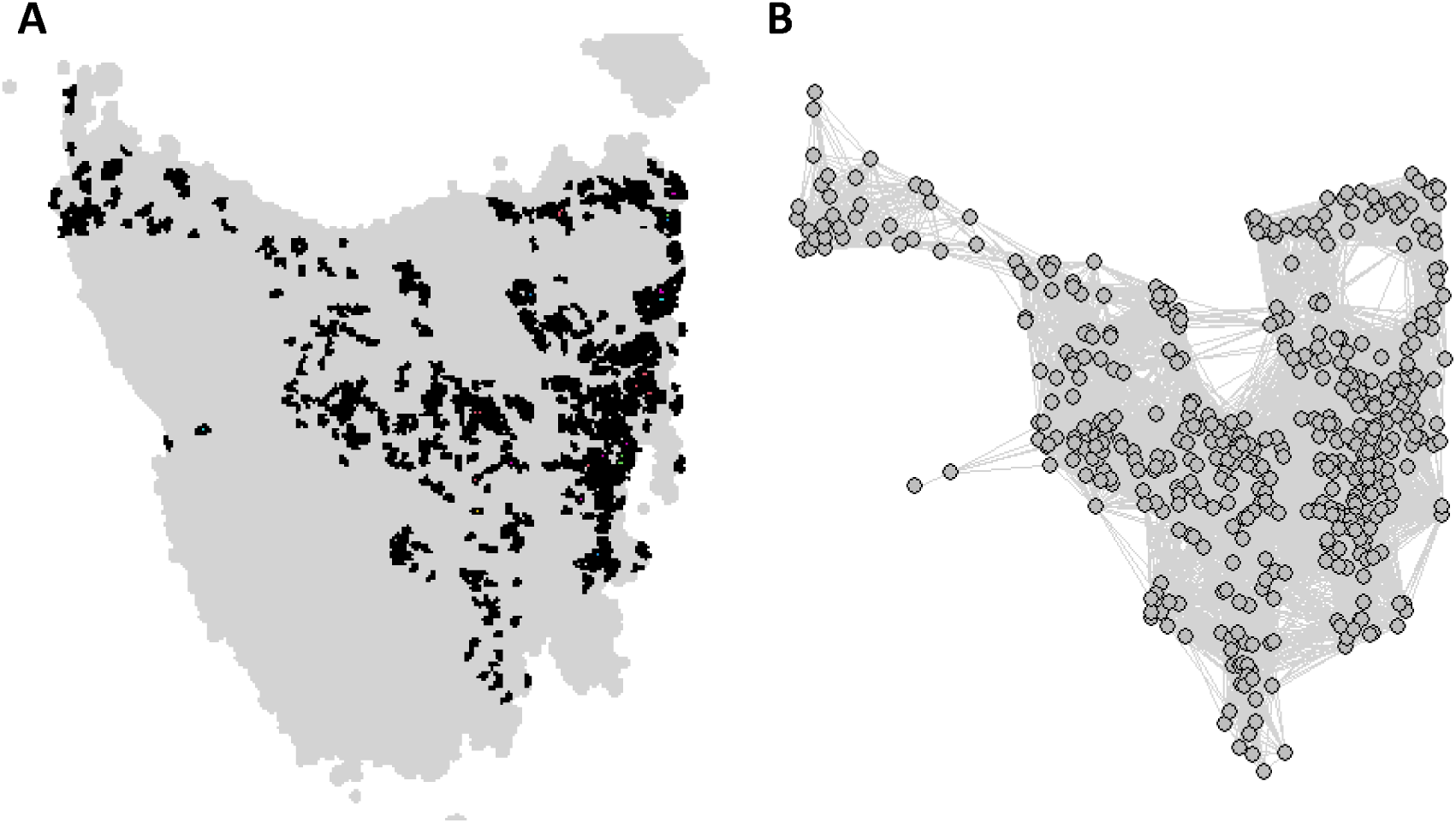
Tasmanian devil metapopulation network map and corresponding habitat. (A) Distribution of sclerophyll forest and coastal heath patches across Tasmania obtained from the TASVEG2 dataset. Patches with an area size <5 km^2^ and those with estimated population density <0.5 devils/km^2^ were excluded. (B) The resulting metapopulation network constructed from A. Each node in the network represents a local population of devils, and each edge (i.e. a connection between two nodes) represents a dispersal corridor between the connected populations.

Dispersal corridors between populations (i.e. the edges in the metapopulation network) were defined according to their relative geographical location. Any two populations within 50 km of each other were assumed to be connected, representing a balance between the largest dispersal events observed in devils (∼110 km) and average estimates based on genetic and demographic analyses (∼14-30 km) (Lachish *et al*. 2011).

### Individual-based model

Each local population in the metapopulation was comprised by a set of individual devils characterised by the following attributes: sex, age, infection status, tumour volume and location within the local population (x and y coordinates). During the course of model simulations individuals were subjected to the following demographic and epidemiological processes in each modelled weekly time step (the time scale of the model) as in (Wells *et al*. 2019):

#### Reproduction

Once a year (i.e. 1 every 52 weeks), female individuals of breeding age (i.e. ≥ 52 weeks old) were randomly selected with a probability of reproducing per breeding season that was determined based on their age (see *Reproduction Probability* in Table 1 for specific probabilities across age groups). Females were not able to reproduce if there were no adult males present within the same local population. Number of offspring recruited as free-roaming individuals from reproducing females was drawn from a binomial distribution (number of events = 4, probability of success = 0.72) according to empirical observations of pouch young survival rates (Guiler 1970a). Each offspring released into the free-roaming population was assigned an age between 32 and 36 weeks with equal probability to account for the duration of the yearly breeding season. This way of simulating the breeding season allowed for computational efficiency (i.e. only one reproductive event per year) while yielding realistic estimates of breeding season duration (i.e. heterogeneity in the aging of offspring released into the population). Furthermore, this approach does not impact disease dynamics as the behaviour of pouch and den young is assumed to be irrelevant to the spread of DFTD. Each offspring was assigned coordinates within their local population equal to those of their mother, sex assigned randomly with equal probability, and a genotype only for population isolation simulations (see “*Tracing the mixing of individuals from different populations*”).

**Table 1:**
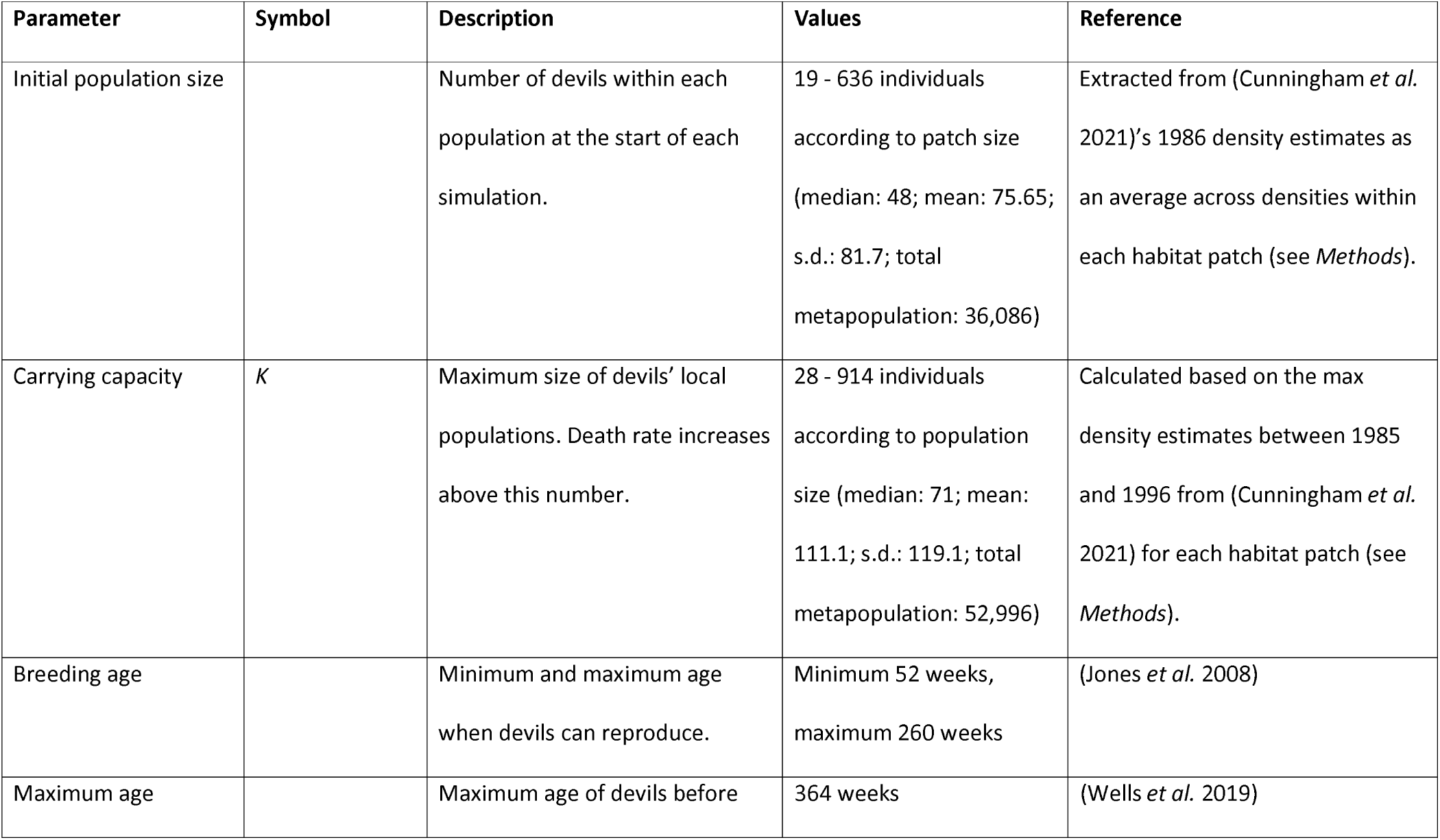
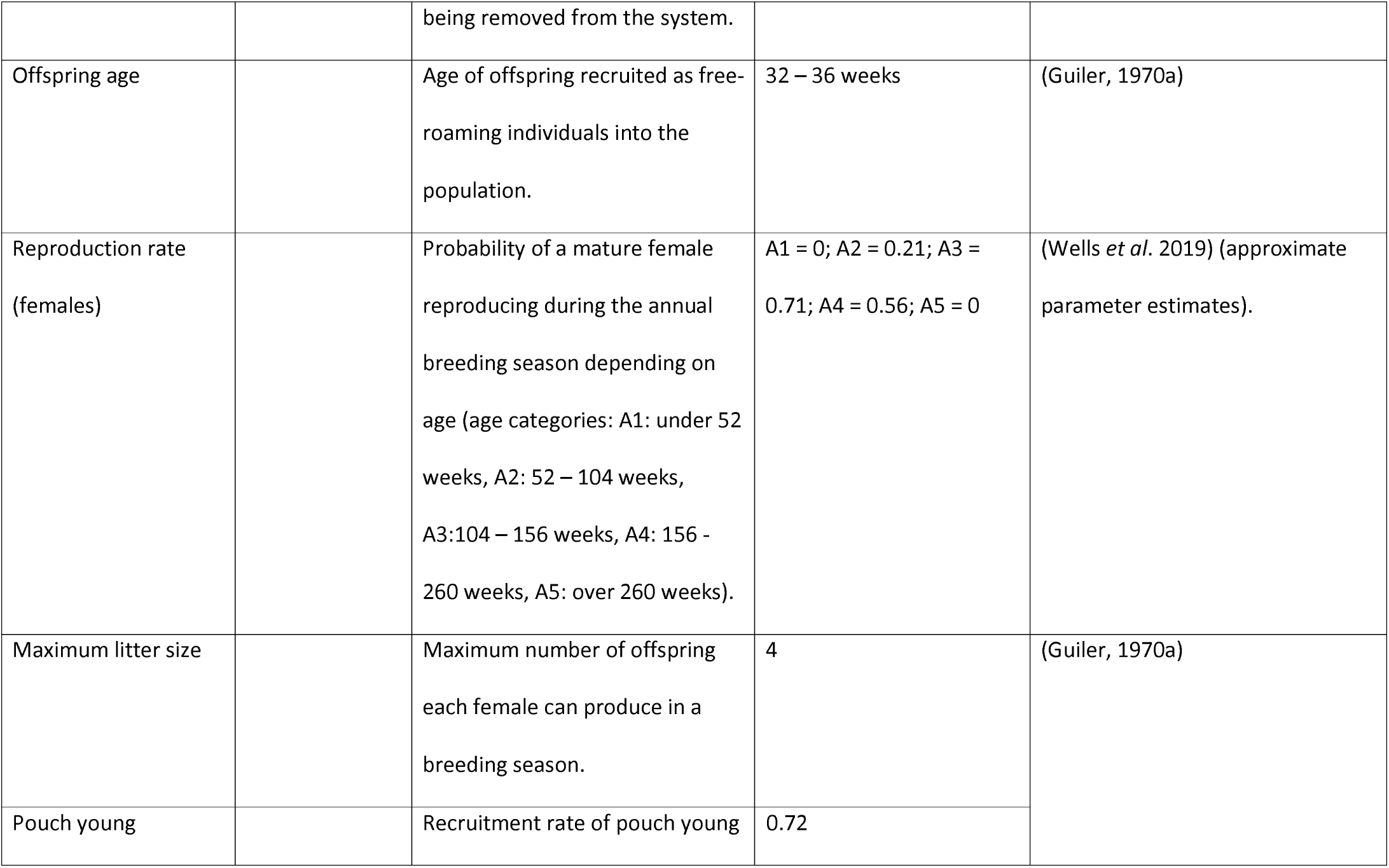
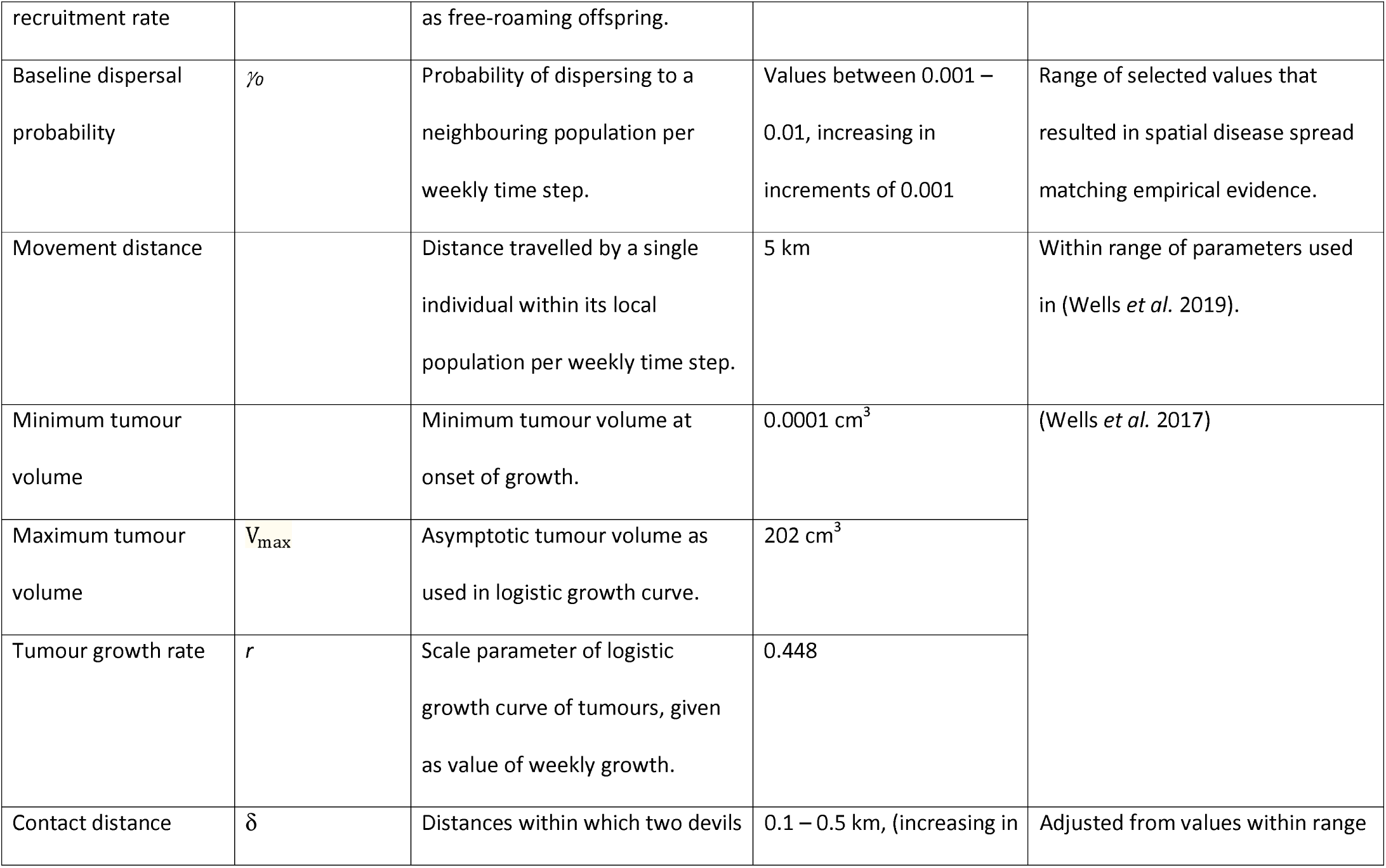
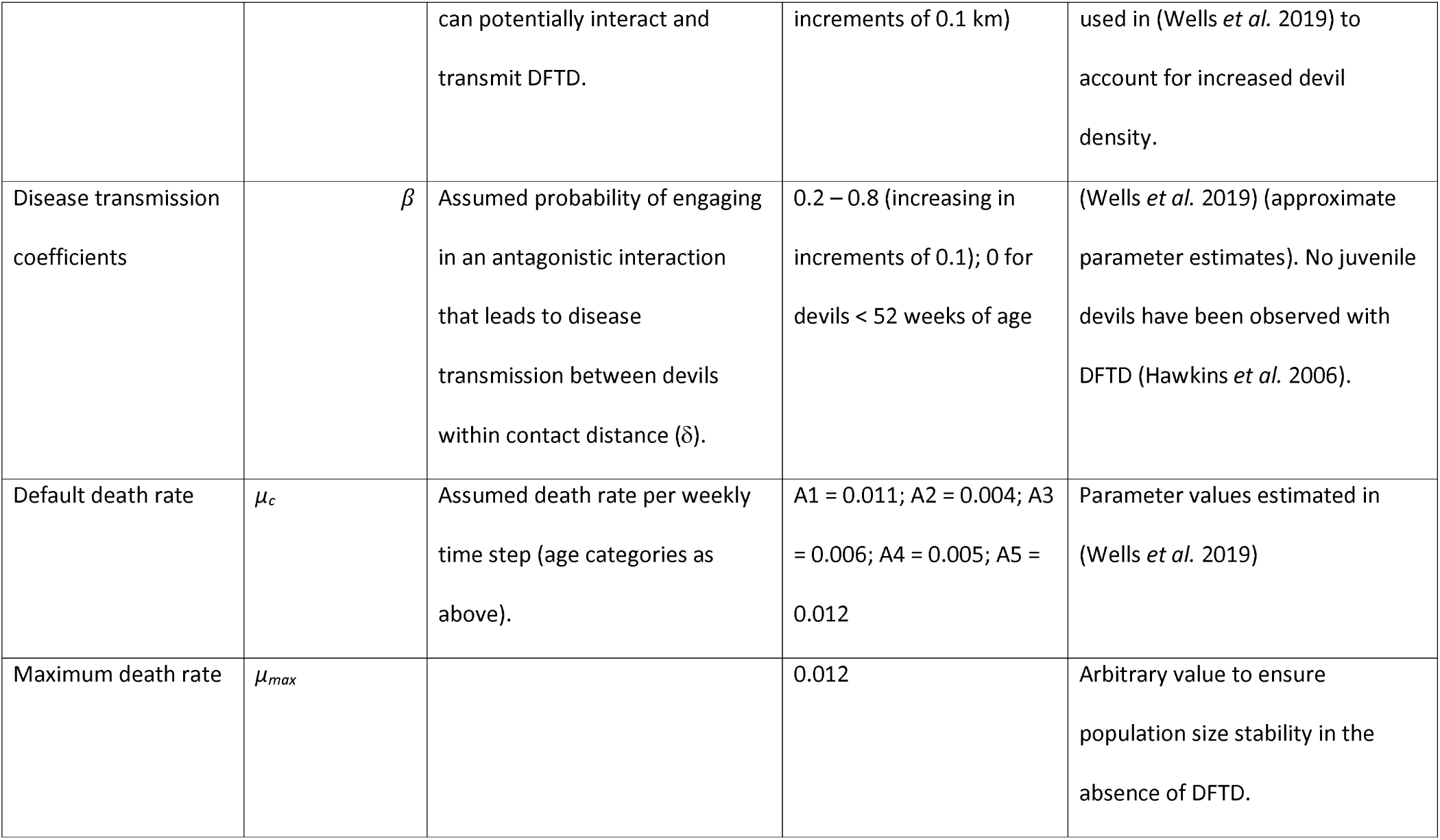
Description and values of the parameters used in the individual-based metapopulation model of Tasmanian devil DFTD spread.

#### Local movement

Movement direction and distance of individual devils within their local population were randomly determined each time step. Direction was randomly sampled from a normal distribution between 0 and 2*π, whereas distance was normally distributed around the average movement speed (mean = 5 kilometres per week, s.d. 0.5). If their movement distance in one direction exceeded the boundaries of the spatial area of the local population, the devil would stop at this boundary. DFTD transmission between individuals was influenced by the maximum pairwise Euclidean distance over which devils were assumed to interact within the weekly time widow (hereafter referred to as the inter-individual contact distance, δ). Larger contact distances would translate into higher levels of within-population mixing, a measure relevant for disease spread.

#### Dispersal

Individuals disperse to neighbouring populations (i.e. connected via a link in the metapopulation) with a certain baseline density-dependent dispersal probability (*γ*_*0*_; Table 1) and proportional to population size with its maximum at carrying capacity thus:

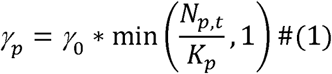

where *γ*_*p*_ is the dispersal probability for devils within population *P, γ*_*0*_ is the baseline dispersal probability, *N*_*p,t*_ is the population size within population *p* at time *t*, and *K*_*p*_ is the carrying capacity of population *p*. Due to increased rates of dispersal being observed in juvenile devils (under 52 weeks old; Lachish *et al*. 2011), these individuals were assumed to have a dispersal probability 10 times higher than adults. Dispersal values used are estimates based on the limited empirical evidence available (Lachish *et al*. 2011) and initial testing in simulations to produce a range of rates of regional disease spread that encompass the observed rate of DFTD spread across Tasmania.

#### Aging

Each time step of the model simulation, one week was added to every individual’s age.

#### Non-DFTD deaths

Population growth was limited by the local populations’ carrying capacity (*K*). Carrying capacity of each local population was calculated using devil density estimates from (Cunningham *et al*. 2021). For each year between 1985 and 1996 (the emergence of DFTD), the TASVEG2 vegetation map was overlaid over Cunningham *et al*.’s density estimates maps and the mean devil density over the area of each habitat patch was extracted. The largest density estimate per patch across these years was used to calculate the corresponding population’s carrying capacity. Local carrying capacities were uniformly scaled to reach a metapopulation-level carrying capacity of 53,000 individuals (the pre-DFTD maximum population size estimates (Cunningham *et al*. 2021) (Supp. Fig. S1B).

Individual death rate (*µ*) was set to a constant when populations were below *K* (*µ*_*c*_, Table 1), increasing with local abundance as the number of individuals in the population exceeded *K*, up to a maximum death rate (*µ*_*max*_) of 0.012 (the maximum death rate value that results in a stable metapopulation size in the absence of DFTD):

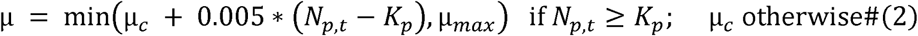

where *µ*_*c*_ is the default death rate for the individual’s age category (see Table 1 for values). Additionally, individuals that reached the maximum age of 364 weeks were considered dead and removed from the system.

#### Infection

The force of infection (*λ*_*i*_) is the probability of an individual becoming newly infected with DFTD in a given week, and is defined as:

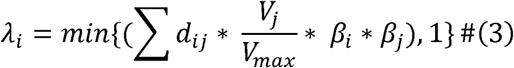

where *d*_*ij*_ is a Boolean indicator of whether a potentially infective individual *j* is within the contact distance (δ) of the susceptible individual *i, V*_*j*_ is the tumour volume of individual *j, V*_*max*_ is the maximum possible tumour volume, and the disease transmission coefficients *β*_*i*_ and *β*_*j*_ represent the probability of antagonistic interaction leading to disease transmission based on individual *i* and *j*’s age. If the susceptible individual becomes infected during an interaction, it was seeded with a tumour of volume 0.0001 cm^3^ (the minimum tumour volume at the onset of growth).

#### Tumour growth

Increase in tumour volume for each individual per week was determined using a logistic function, adapted from Hamede *et al*. (2017):

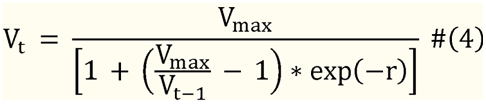

where *V*_*t*_ is tumour volume in the current weekly time step, V_t-1_ is the tumour volume in the previous time step, V_max_ is the maximum tumour volume, and *r* is tumour growth rate in cm^3^ per week (Table 1).

#### DFTD-induced mortality

Even though data on devils’ causes of mortality is sparse, previous studies have found that DFTD-induced mortality is close to 100%, with animals succumbing to the disease within 6 to 18 months (Hamede *et al*. 2012; Wells *et al*. 2019). For simplicity, we assume a 100% mortality rate. Using individual-based simulated mortality data from our model we estimated the tumour volume-dependant mortality rate to ensure a mean infection-to-death time period of 35 weeks (well within the estimates from the empirical data) (Supp. Fig. S2). Thus, disease-induced mortality Ω_size_ per time step was assumed to be proportional to the individual’s tumour volume and given by:

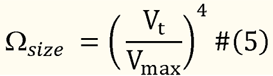

If an individual was selected to die in that time step, it was removed from the population.

### Tracing the mixing of individuals from different populations

To assess the extent to which metapopulation fragmentation (via population isolation) could potentially restrict gene flow between local devil populations, we traced between-population genetic mixing as a proxy for devil’s genetic diversity across the metapopulation. Individuals were assigned a population-specific “genotype” at the beginning of each simulation. Genetic inheritance during reproduction followed a simple model in which all adult males in a population have an equal probability of fathering local offspring. All offspring from a single brood were assumed to be fathered by the same male individual. For simplicity offspring inherited either the mother or father’s genotype with equal probability. Population mixing was measured for each local population using Nei’s within population variation index *H*_*i*_ (Nei 1973):

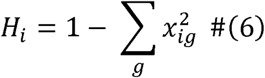

where *x*_*ig*_ is the proportion of individuals in population *i* with the genotype *g*. When a population was empty, its genetic variation was set to zero. This measure of population mixing was only used in habitat fragmentation experiments (see below).

### Model Simulations

During each time step (1 week), the following events occurred in order: (1) dispersal of individuals between local populations in the metapopulation network, (2) movement of individuals within populations, (3) offspring release into the free-ranging population (once every 52 timesteps only), (4) death due to DFTD, (5) tumour growth, (6) disease transmission between individuals within specified contact distance, (7) non-DFTD deaths, and (8) aging.

At the start of each simulation, local population size (i.e. number of individuals) was initialised independently for each population using population density estimates for the year 1986 (Cunningham *et al*. 2021), habitat patch area, and the same density multiplier used to calculate carrying capacity (see *Metapopulation structure*; Table 1). Age, sex and location (i.e. coordinates in space) of each individual were randomly assigned. Age was uniformly sampled between 32 and 364 weeks; sex was randomly assigned with .5 probability; and spatial coordinates (x, and y independently) were uniformly sampled between 0 to the square root of the population’s habitat patch area (i.e. the habitat patch was assumed to be square shaped). After a 520 week (i.e. ten years) burn-in period in the absence of disease to ensure the stability of metapopulation dynamics and demographics before the disease introduction, three randomly chosen individual devils from the population closest to the location where DFTD was first recorded in Tasmania (Hawkins *et al*. 2006) were infected with DFTD to minimise the risk of all initially infected individuals dying before the outbreak is established.

To investigate the effect of varying contact distance (δ), disease transmission coefficients (*β*), and the baseline dispersal rate (*γ*_*0*_) on population size, disease prevalence and population stability, we ran the model described above for a total of 1820 weekly time steps (35 years) for each combination of these parameters’ values (Table 1). Contact distance took values from 0.1 to 0.5 kilometres, with increments of 0.1 km; disease transmission coefficients were varied between 0.2 – 0.8 increasing by 0.1; and baseline dispersal rate values ranged from 0.001 to 0.01, with increments of 0.001 (a dispersal probability of 0.001 is equivalent to 520 dispersal events per 10,000 individuals per year). We performed 20 replicated simulations for each parameter combination, resulting in 7,000 simulations in total.

### Metapopulation analysis

Metapopulation dynamics and disease spread resulting from the parameter combinations introduced above were assessed using two summary measures:

1. Total metapopulation size: quantified as the median, across the last 520 weeks (ten years) of the simulation, of the sum of the sizes (i.e. number of individuals) of all local populations at each time step.
2. Proportion of local populations where at least one case of DFTD was recorded during the entire simulated time period.

### Pattern matching of disease spread to empirical data

We identified the parameter combination from our simulations for which the spatial patterns of disease emergence most closely matched a recently published empirical predictions of the disease front from 1996 to 2015 (Cunningham *et al*. 2019) and predicted metapopulation size declines from 1986 to 2020 (Cunningham *et al*. 2021). Parameter combinations were first filtered to match estimates of metapopulation size for the year 2020 from empirical data according to the 95% credible intervals of estimates reported by Cunningham *et al*.’s (2021) and with a DFTD prevalence ≥ 5%. From the remaining parameter combinations, the one with the best match for disease arrival at different locations in Tasmania through time (assessed within five-year time windows of empirical estimates from Cunningham *et al*.’s (2019)) was assumed to represent the most realistic simulation scenario. Matching of disease arrival time between simulated and empirical data was quantified as the proportion of local populations falling within the correct temporal disease front as reported by Cunningham *et al*.’s (2019). The parameter combination thus selected was used for the metapopulation fragmentation experiments.

### Metapopulation fragmentation experiments

To evaluate the consequences of population isolation / metapopulation fragmentation on population dynamics and disease spread, we ran a series of simulation experiments where up to 100 local populations were isolated from the metapopulation. Between 10 and 100 populations were isolated (incrementally in sets of 10; i.e. 10, 20, 30 … 100) at week 1040 of each fragmentation simulation. We implemented three different isolation methods based on the connectivity profiles of local populations:

1. Random: Populations to be isolated were randomly chosen.
2. Degree: Isolated populations were chosen in order according to their degree (i.e. the number of connections of the local population), from the most connected to the least connected population.
3. Betweenness: Populations to isolate were selected in order according to their betweenness centrality, a node-level network measure that quantifies the extent to which the shortest paths connecting any two nodes (populations) in the network (the metapopulation) comprise the focal node. Local populations were thus ranked from most to least central according to their betweenness centrality, and those with the highest centrality scores were removed first. Betweenness centrality of a local population *v* was calculated thus (Freeman 1979):

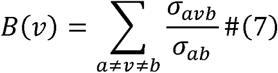

where *σ*_*ab*_ is the number of shortest paths that between population *a* and population *b*, and *σ*_*avb*_ is the number of these that pass through population *v*.

We performed 30 replicated simulations for each isolation method and number of populations isolated, with DFTD being introduced 520 weeks (10 years) into the 1820-week (35 year) simulation. To compare the outcomes of the disease dynamics to the neutral (i.e. no disease) scenario, we performed further 20 replicated simulations without DFTD being introduced. This resulted in 1,500 simulations in total. Total metapopulation size, disease prevalence, and proportion of local populations reached by DFTD were calculated as above. Additionally, the mean within-population genetic variation (Eq. 6) was quantified to assess the effects of fragmentation on genetic composition of the metapopulation.

Model implementation and analyses were carried out using R version 4.1 (R Core Team 2020). Source code for the full model and scripts used for data analyses is available from the GitHub repository: https://github.com/RowanDurrant/DFTD.

## Results

### Disease spread is driven by the interplay between local transmission and regional movement

Population size decreased as dispersal rate (*γ*_*0*_) increased, across all values of transmission rate (*β*) and contact distance (δ). However, only under scenarios of relatively large transmission rate (*β* >= 0.5) did increasing contact distance and dispersal rate lead to considerable reductions in overall metapopulation size (Fig. 2). This suggests that all three factors interact synergistically to cause major population declines.

**Figure 2.**
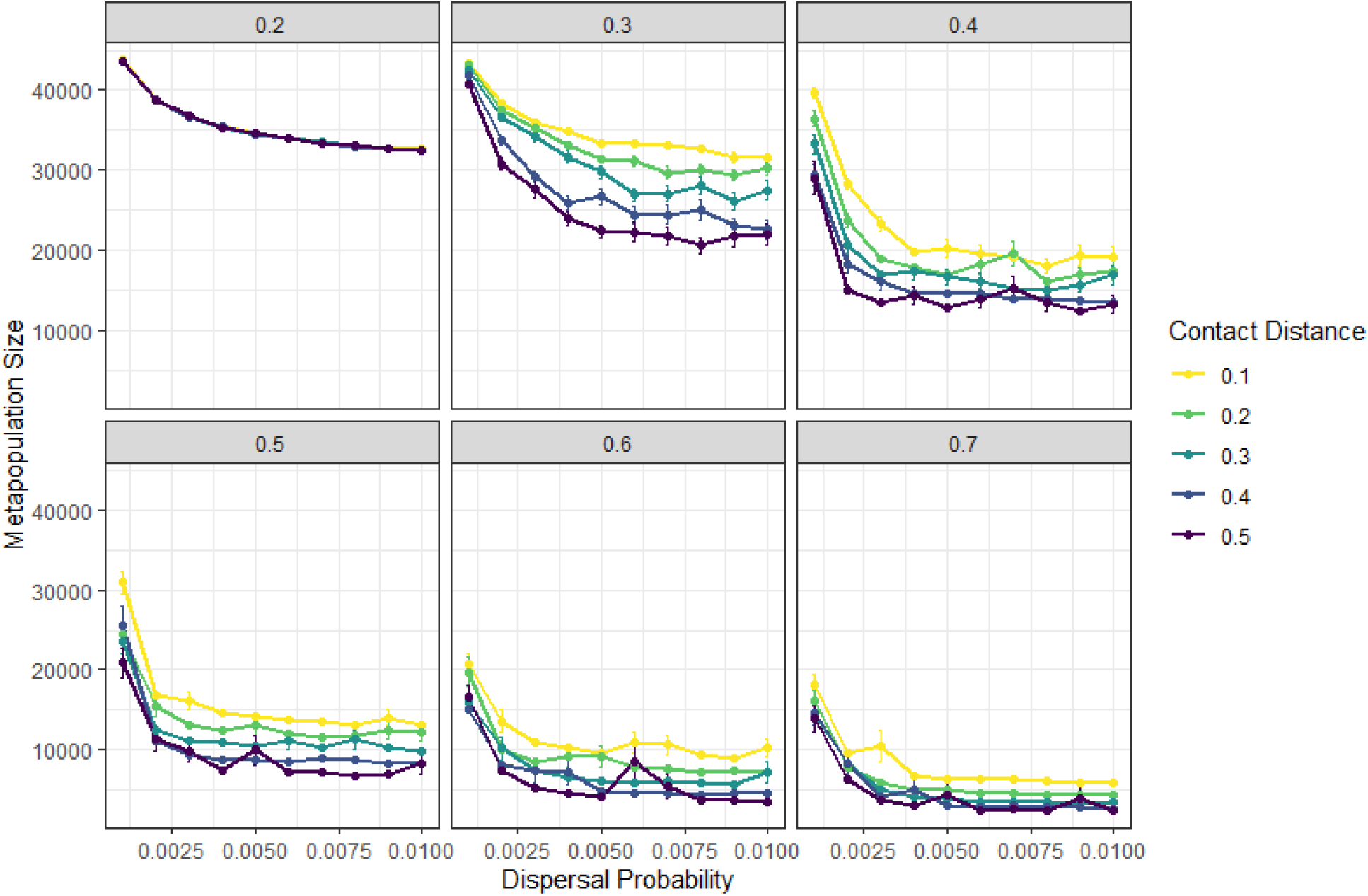
Tasmanian devil metapopulation size is influenced synergistically by within-population mixing (contact distance), dispersal rate and transmission probabilities. Points and vertical bars show the mean and standard error (across 20 replicated simulations) of median values of metapopulation size taken over 520 weeks (10 years) of each simulation. Panels display results for different transmission probability scenarios (numbers at the top of the panel). Colour of points and lines represent different contact distance scenarios.

At relatively low transmission probability values (e.g. *β*= 0.2), metapopulation size declined as dispersal rate increased at the same rate across all values of contact distance. This pattern was also observed in the absence of disease (Supp. Fig. S3), suggesting that this population decline is caused by source-sink metapopulation dynamics independent of disease effects.

At transmission probability of 0.2, DFTD failed to spread far beyond the initially infected population, across all values of dispersal rate and contact distance (Fig. 3). At higher values of disease transmission, however, the proportion of infected populations rises with the increase in dispersal probability. Contact distance only has an effect on the proportion of populations reached by DFTD at a transmission probability of 0.3, with a negligible influence at higher values (Fig. 3).

**Figure 3:**
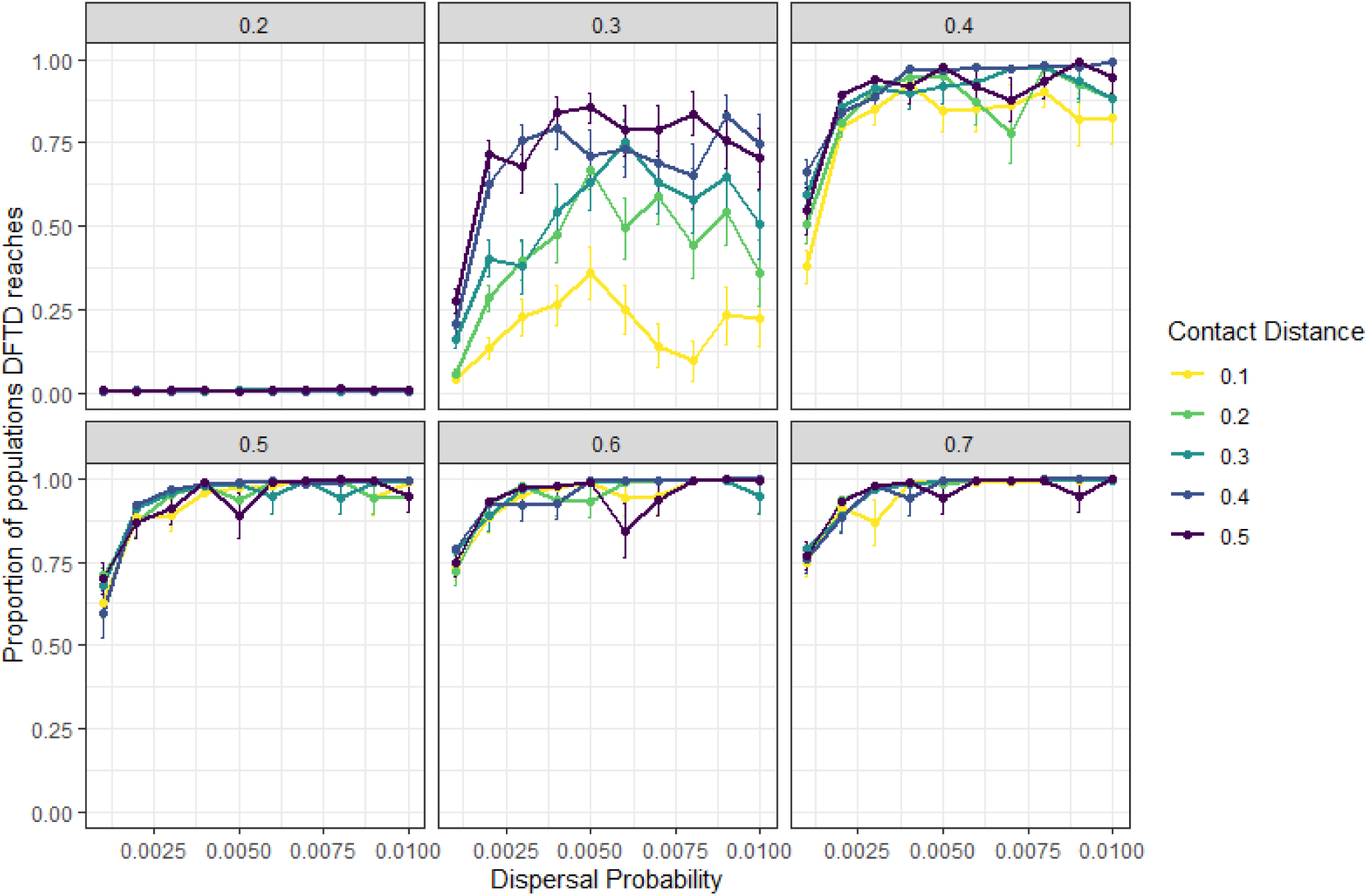
Average proportion of local devil populations reached by DFTD is influenced by contact distance, dispersal and transmission probabilities. Points and vertical bars show the means and standard error of median values of the fraction of local populations within the metapopulation reached by disease within the time period of the simulation, taken over the last 520 weeks (10 years) of each simulation. Panels display results for each value of transmission probability (numbers at the top of each panel). Point and line colours represent different values of contact distance.

### Spatial spread of DFTD across Tasmania is explained by metapopulation dynamics

We compared our simulation results to recent empirical DFTD disease front predictions in order to understand the metapopulation mechanisms that might be behind DFTD spread. Out of all parameter combinations explored, the scenario that most closely matched the observed DFTD disease front (Fig. 4B) was that of contact distance = 0.1, baseline dispersal rate = 0.009 and transmission probability = 0.4. In this scenario, 67.5% of populations fell within the correct disease arrival wave (i.e. matched the year of arrival reported in previous research (Cunningham *et al*. 2019)). The mean year of arrival of DFTD at each local population from simulations using these parameter values resembled empirical observations, only reaching the north-western corner of Tasmania within the last years of the simulation (Fig. 4A).

**Figure 4.**
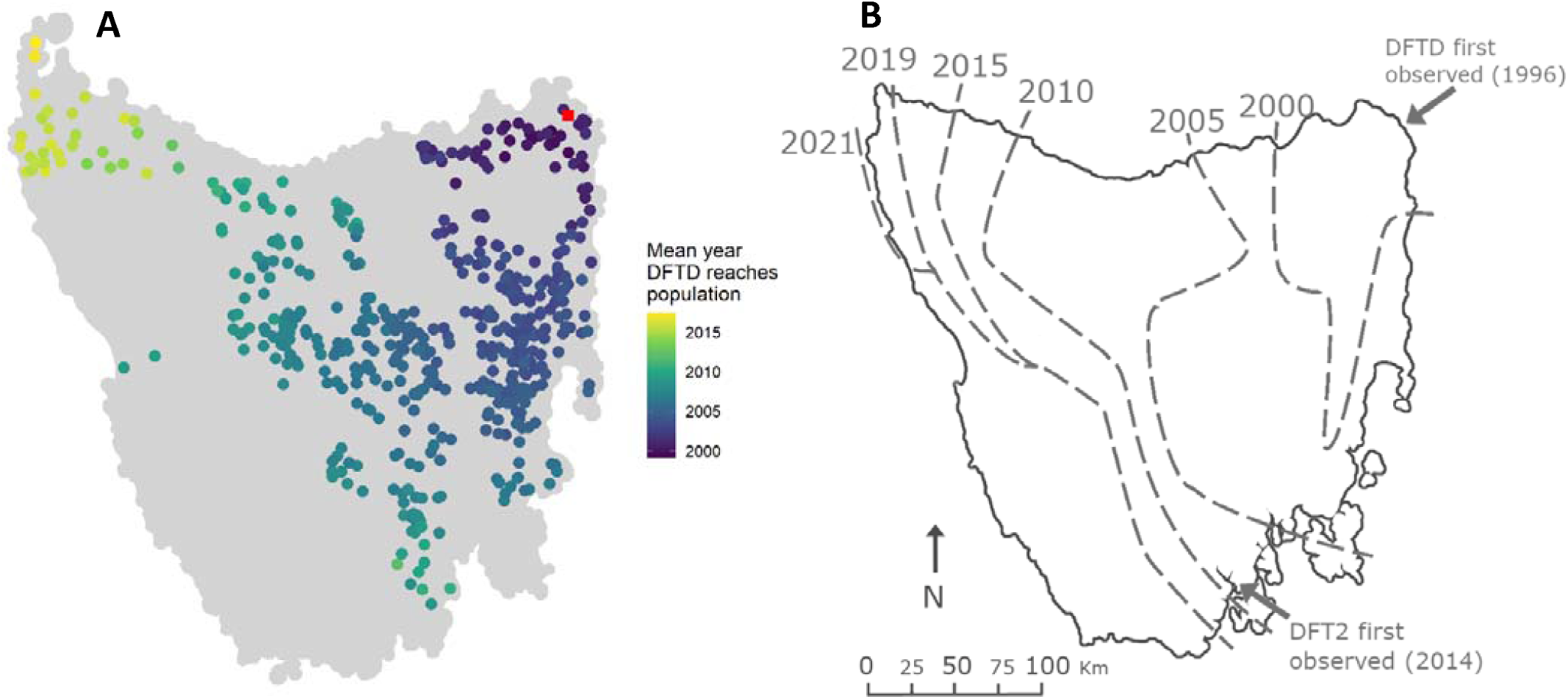
DFTD arrival wave across Tasmania is well approximated by the mechanistic metapopulation model. (A) Year of DFTD arrival at each local population based on averaged outputs from metapopulation simulations using the parameter combination that best matches empirical data (δ = 0.1, _*0*_ = 0.009 and = 0.4). The initially infected population at the beginning of simulations is represented by a red square. Point colour represents the year of DFTD arrival; grey points are populations that DFTD did not reach within 25 years of disease introduction. (B) Map of the estimated disease front in Tasmania based on empirical data analysis (modified from (Cunningham *et al*. 2019)). The map represents the data used to match simulation results and identify the parameter combination used to obtain results presented in (A).

Metapopulation size and disease prevalence of simulations with this parameter combination exhibit a strong decline and increase respectively up until approximately 700 weeks after DFTD introduction (∼ week 1200), with signs of a metapopulation recovery after that (Fig. 5). The final increase in cumulative populations infected after the brief plateau corresponds to the north-western populations which only become infected towards the end of the simulation, with the disease front progressing more slowly than in previous years. These patterns resemble those observed from empirical data during comparable time periods (Fig. 5 in Cunningham *et al*. 2021).

**Figure 5.**
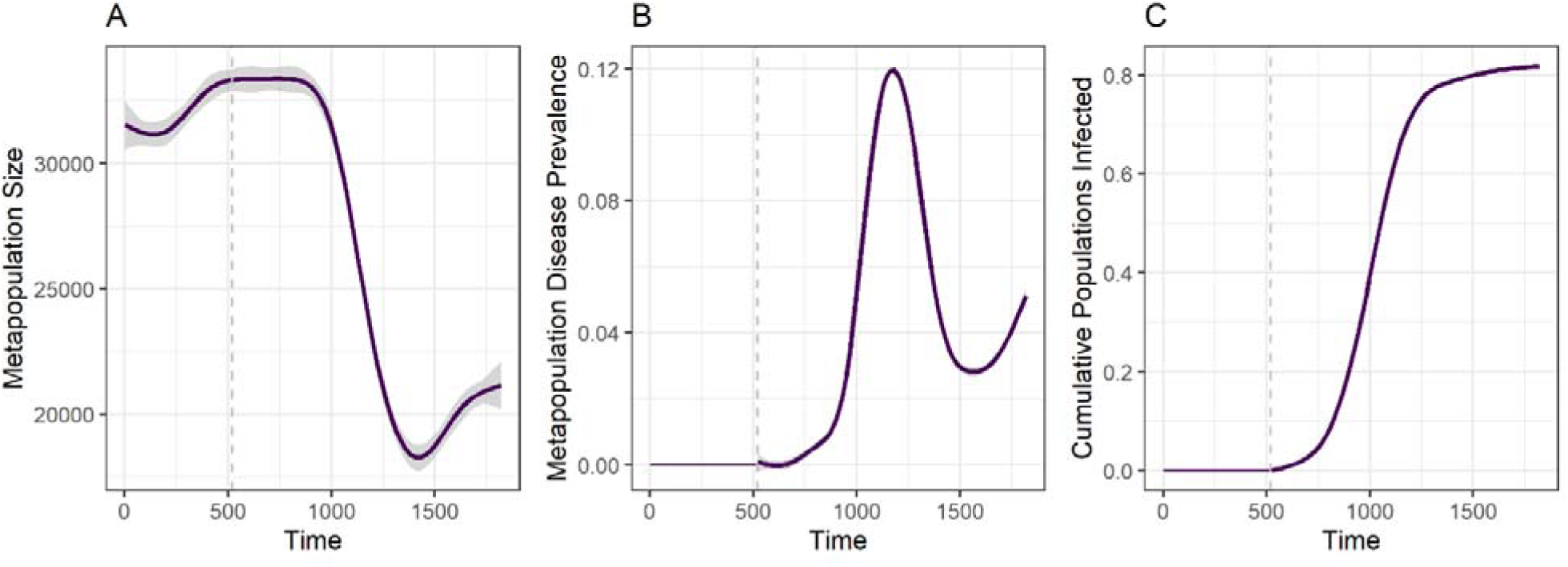
Metapopulation dynamics of disease spread matching the empirically observed disease spread front. Total metapopulation size (A), metapopulation-level DFTD prevalence (B), and cumulative fraction of populations reached by DFTD (i.e. have had at least one diseased individual during the course of the simulation) (C) through time. Vertical grey dashed lines indicate time of DFTD introduction. Lines and shadows around them represent mean and standard error values across 20 replicated simulations with parameters δ = 0.1, *γ*_*0*_ = 0.009 and *β* = 0.4.

### Fragmentation reduces disease spread at the cost of population size

Metapopulation fragmentation experiments resulted in similar magnitudes of total metapopulation size decline with increasing numbers of populations being isolated regardless of whether populations were isolated at random or according to their position in the metapopulation network (degree or centrality) (Fig. 6A). The proportion of populations reached by DFTD varied between population isolation methods. Isolation according to betweenness centrality or at random resulted in declines in the number of populations with DFTD presence as the number of isolated populations increased. Population isolation by degree (i.e. number of connected neighbouring populations) did not show such relationship (Fig. 6B).

**Figure 6.**
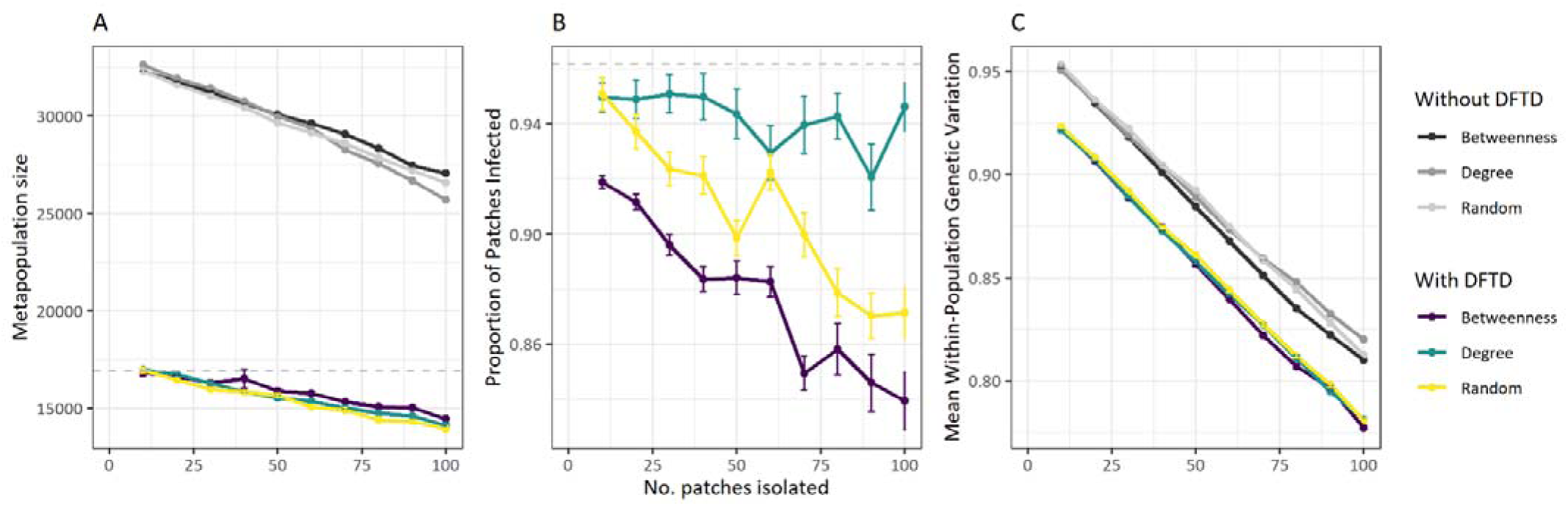
Effects of fragmentation on DFTD spread in Tasmanian devil metapopulations. Metapopulation size (A), proportion of individuals infected within the metapopulation (B) and mean within-population genetic diversity index (C) are influenced by the number of populations isolated from the metapopulation. Proportion of populations infected is also influenced by whether populations are isolated based on the population’s centrality in the metapopulation (i.e. betweenness), its connectivity (i.e. degree) or at random (B). Colours represents different population isolation methods. Points and vertical bars display the average and standard error of the different quantities across 20 replicated simulations using the parameter combination that best matches the spatial wave of disease spread (δ = 0.1, *γ*_*0*_ = 0.009 and *β* = 0.4; Fig. 4).

As expected, within-population genetic diversity also decreased as the number of populations isolated increased for all population isolation methods (Fig. 6C). These patterns were also observed in simulations without DFTD, although both metapopulation size and population genetic diversity were considerably higher in the absence of DFTD, illustrating that population isolation does not curtail the negative impact of DFTD at the metapopulation level (Fig. 6).

## Discussion

We developed an individual-based metapopulation model that allowed us to investigate the synergistic effects of local-scale disease transmission processes and population dynamics (e.g. transmission rate and inter-individual contact distance) and landscape-scale metapopulation processes (e.g. dispersal) on the spread of a transmissible cancer in Tasmanian devil populations, with the consequences for population abundances that ensue. We matched the outcomes of this model to published empirical data and identified parameter combinations able to reproduce observed patterns of disease spread. We further used this information to perform *in-silico* metapopulation fragmentation experiments to investigate the potential effects of population isolation measures on DFTD spread and its consequences for population persistence and dynamics. Our results suggest that inter-individual contact distance -a measure of within-population mixing and contact facilitating disease transmission-, dispersal of individuals among local populations, and DFTD transmission rates interact in non-intuitive ways to drive population declines in Tasmanian devils.

Whereas dispersal was identified as the main driver of regional DFTD spread, this was only observed for relatively high DFTD transmission rates (≥ 0.3). The relevance of inter-individual transmission for effective spread is reinforced by our results for intermediate transmission rates (0.3 and to some extent 0.4). These levels of transmission alone are still not high enough to ensure complete spread, unless contact distance between individuals is high enough to increase transmission events. These observations hint at the critical role of within-population mixing and transmission rate in determining DFTD prevalence, and associated individual death, at the local population level. This would eventually translate into an overflow of disease into neighbouring populations, creating a source-sink phenomena, and ultimately resulting in regional disease spread.

The implication that local within-population mixing and dispersal probability both influence the magnitude of a regional outbreak means that a management method that targets either one could prevent a local outbreak from progressing. We hypothesised that isolating some local populations would reduce disease spread, resulting in a larger population size, but at the risk of decreasing population mixing and the potential associated loss of genetic diversity. While we predicted correctly that population mixing would decrease leading to lower within-population genetic diversity, population size also declined as the number of isolated patches increased, and the proportion of populations that DFTD reached varied depending on the population isolation method. Isolating populations with the highest betweenness was the most effective approach for reducing metapopulation DFTD prevalence, which is in line with previous suggestions that immunising nodes in a network with high betweenness is more effective than immunising those with a high degree (Yang & Wang 2016). Furthermore, isolating populations with a high degree in our simulations was less effective at reducing DFTD prevalence than isolating at random, which was counter to both our expectations and the results of previous immunisation studies (Pastor-Satorras & Vespignani 2002). One explanation as to why isolating populations by their number of connections to neighbouring populations (i.e. degree) had little effect on the spread of DFTD across the metapopulation is that the 100 populations with the highest degree were all clustered in one geographic area where DFTD was already present before the populations were isolated, and so their isolation did not lie in the way of the disease front. In other metapopulation structures where this clustering pattern is not seen, we would expect isolation by degree to have a greater influence on the course of the outbreak, as observed by Pastor-Satorras & Vespignani (2002). The place of introduction of an emerging disease, landscape context and also the timing of any spatial intervention strategy in relation to previous disease spread may therefore determine which spatial isolation methods may be most efficient.

We found that the number of devils living in populations that were isolated declined rapidly, reducing to zero in some cases (Supp. Fig. S4). Due to their isolation, these populations were unable to be recolonised by individuals from neighbouring populations, suppressing thus the ‘rescue effect’ commonly observed in metapopulation dynamics (Brown & Kodric-Brown 1977; Gotelli 1991) and highlighting its importance for devil population persistence. On the other hand, populations that remained within the metapopulation network were quickly recolonised if their population size dropped to zero (Supp. Fig. S5). As we increased the number of isolated populations, more of these non-recolonisable population extinctions occurred. It has long been argued whether increased connectivity between populations is beneficial or harmful during a disease outbreak, with some arguing that wildlife corridors could facilitate the spread of disease (Hess, 1994), and other studies showing that local populations benefit from recolonization by individuals from neighbouring populations (Levins 1969; Jousimo *et al*. 2014). In our simulations, while DFTD indeed spread further in more highly connected metapopulation structures, isolating populations, suppressing thus recolonisation, caused more harm to the devil population than the potential benefits of reducing disease spread would confer. This result also supports Siska *et al*.’s (2018) suggestion that population recolonisation is an important factor in preventing devil extinction. Together these findings suggest that trying to completely isolate some populations to reduce the long-distance movement of devils would not be a suitable method to prevent further spread of an ongoing regional DFTD outbreak, and that while Hess (1994) was correct in correlating increased metapopulation connectivity with increased disease spread, the benefits of this connectivity far outweighs the risks in this scenario. Population isolation through fencing may be unsuitable for devils as the disease is already present throughout the species distributional range. The only area where the disease has not been detected is southwestern Tasmania’s temperate rainforest and button grass (sub-optimal habitat for the species), where fencing is not logistically feasible. Isolation may be a viable management method when accompanied with efforts to actively maintain the size and genetic diversity of isolated populations, or in species where population size and genetic diversity is less of a concern.

Vaccination has been proven to be effective in reducing disease spread in other wildlife diseases, such as rabies (Brochier *et al*. 1991). The release of captive-bred, immunised devils into local populations has been previously suggested to be a viable option in reducing DFTD spread (Bruno *et al*. 2017). Devils have been injected with sonicated DFTD cells with the aim to stimulate and adaptive immunity as a potential vaccine (Tovar et al 2017), however, there is no evidence that attempted immunizations are prophylactic in the wild (Pye et al 2018; Owen & Siddle 2019). Furthermore, empirical and theoretical evidence suggest that vaccinations that do not prevent transmission and spread of disease (often referred as leaky or imperfect vaccines) can create ecological and epidemiological conditions that would allow more virulent pathogen strains to emerge and persist (Gandon *et al*. 2001; Read *et al*. 2015). In the case of the Tasmanian devils and DFTD, an increasing number of studies have demonstrated natural adaptations to the epidemic, with devils developing defence mechanisms against infection (Hohenlohe *et al*. 2019; Hamede *et al*. 2020a) and genetic changes in the tumour leading to reduced transmission and epidemic outcomes (Hamede *et al*. 2015; Patton *et al*. 2020). Future research should integrate these devil-tumour evolutionary processes to evaluate their long-term effects on disease spread and metapopulation dynamics.

Whilst it is too late to prevent DFTD from spreading across Tasmania, it is hoped that alternative management options can be developed to prevent the spread of a second and independently evolved transmissible cancer affecting Tasmanian devils, devil facial tumour 2 (DFT2) (Pye *et al*. 2016) from causing another wave of catastrophic population decline (James *et al*. 2019; Flies *et al*. 2020). DFT2 is so far confined to the geographic peninsula where it was discovered in 2014, in south-eastern Tasmania (James *et al*. 2019). Although there is currently very little information on the epidemiology and population effects of DFT2, localised spatial spread of DFT2 has been found to be much slower than in DFTD (James et al. 2019). DFTD has spread throughout Tasmania over continuous habitat at a rate of 25 km per year (McCallum *et al*. 2007), however, DFT2 is confined to a peninsula bounded by water on the sides (east, west and south), and a highly urbanised landscape to the north, which has resulted in a spatial spread of 7 km per year (James *et al*. 2019). Another factor contributing to the slow spread of DFT2 might be the current low devil population densities across Tasmania, combined with human modified and fragmented landscapes in the surrounding areas where DFT2 emerged. Future models could integrate DFTD-DFT2 infection dynamics and assess the role of highly urbanised-fragmented landscapes on disease spread, and their effects on metapopulation dynamics.

Counter to empirical observations where disease prevalence is observed to remain high behind the disease front (Lazenby *et al*. 2018), our model predicts a peak in DFTD cases that begins to decline shortly after the devil metapopulation size starts to decline, and rises again when the metapopulation begins to recover. This pattern was also observed by Wells *et al*., (2017). One explanation as to this departure from field observations is that both models use density dependent transmission instead of frequency dependant transmission, and that the existence of a host density threshold could cause this reduction in disease prevalence. As there is strong evidence to suggest frequency-dependence in DFTD transmission (McCallum *et al*. 2009), future versions of our model could incorporate frequency-dependent transmission to investigate whether this can reproduce the observed patterns of DFTD persistence. We also observed the metapopulation size decreasing as dispersal probability increased, even in the absence of DFTD. It is likely that the increased dispersal of individuals results in source-sink dynamics in which well-connected populations act as pseudo-sinks (populations that are net receivers of dispersers but do not necessarily depend on them to remain viable or increase their relative contribution as sources; Watkinson & Sutherland, 1995), as previously observed by Zamberletti *et al*. (2018), and so are pushed above their carrying capacity, resulting in an increased death rate (Supp. Fig. S6).

Our model also does not consider seasonal changes in transmission, social networks of devils, or the presence of DFT2, all of which may be beneficial to include in future models of DFTD. However, such a model could be used in the future to optimise alternative management strategies and forecast future disease dynamics in the devil-DFTD-DFT2 system. Our work highlights the view that host-pathogen theory, population and ecological processes can be applied to reveal the dynamics of infectious cancers in wildlife and further understand how populations and species respond to oncogenic processes.

## Acknowledgements

We acknowledge the support of the Supercomputing Wales project, which is part-funded by the European Regional Development Fund (ERDF) via Welsh Government. The Australian Research Council provided financial support for collecting data to parameterise our models (DE170101116; LP170101105). We also wish to thank Hazel Nichols and Jon Bielby for their useful feedback on an early draft.

## Supplementary Material

**Supplementary Figure S1:**
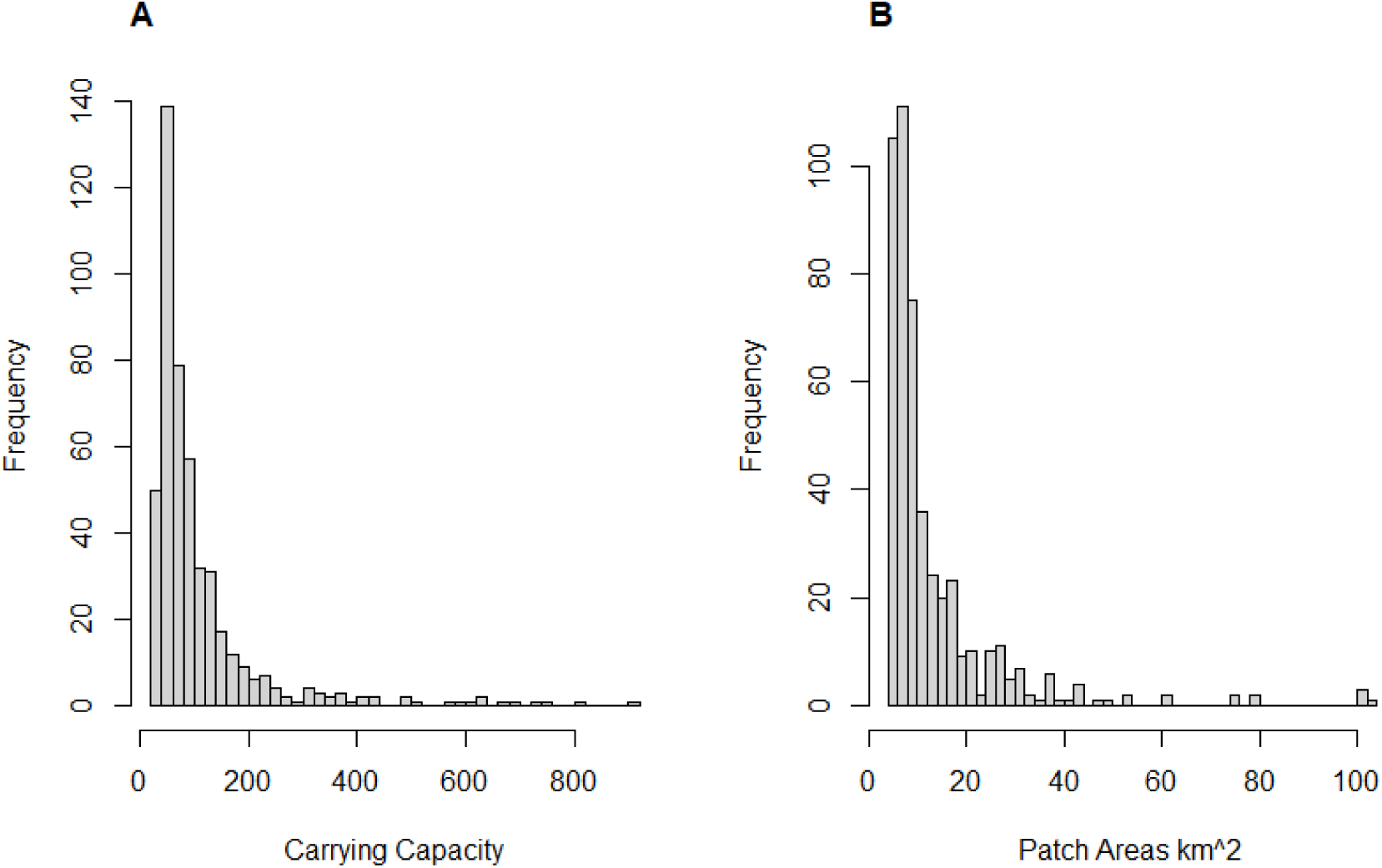
Distribution of carrying capacities and patch area sizes resulting from the calculations of habitat patch areas and population sizes extracted from vegetation maps and density estimates (see *Methods*).

**Supplementary Figure S2:**
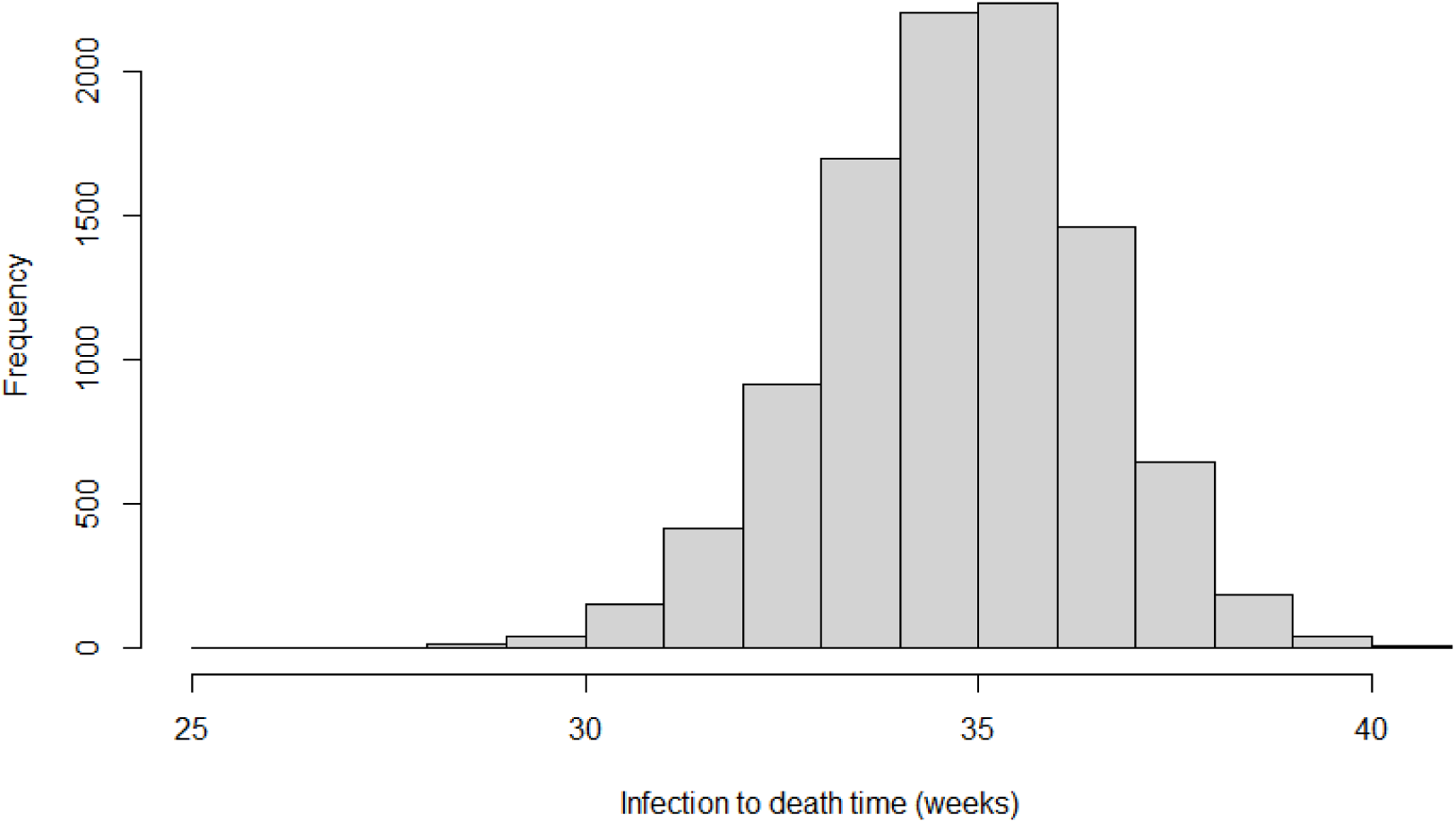
Distribution of infection to death time values (in weeks) for individuals tracked over the course of an entire simulation using equation (Eq. 5) to calculate death rate (see *Methods*).

**Supplementary Figure S3:**
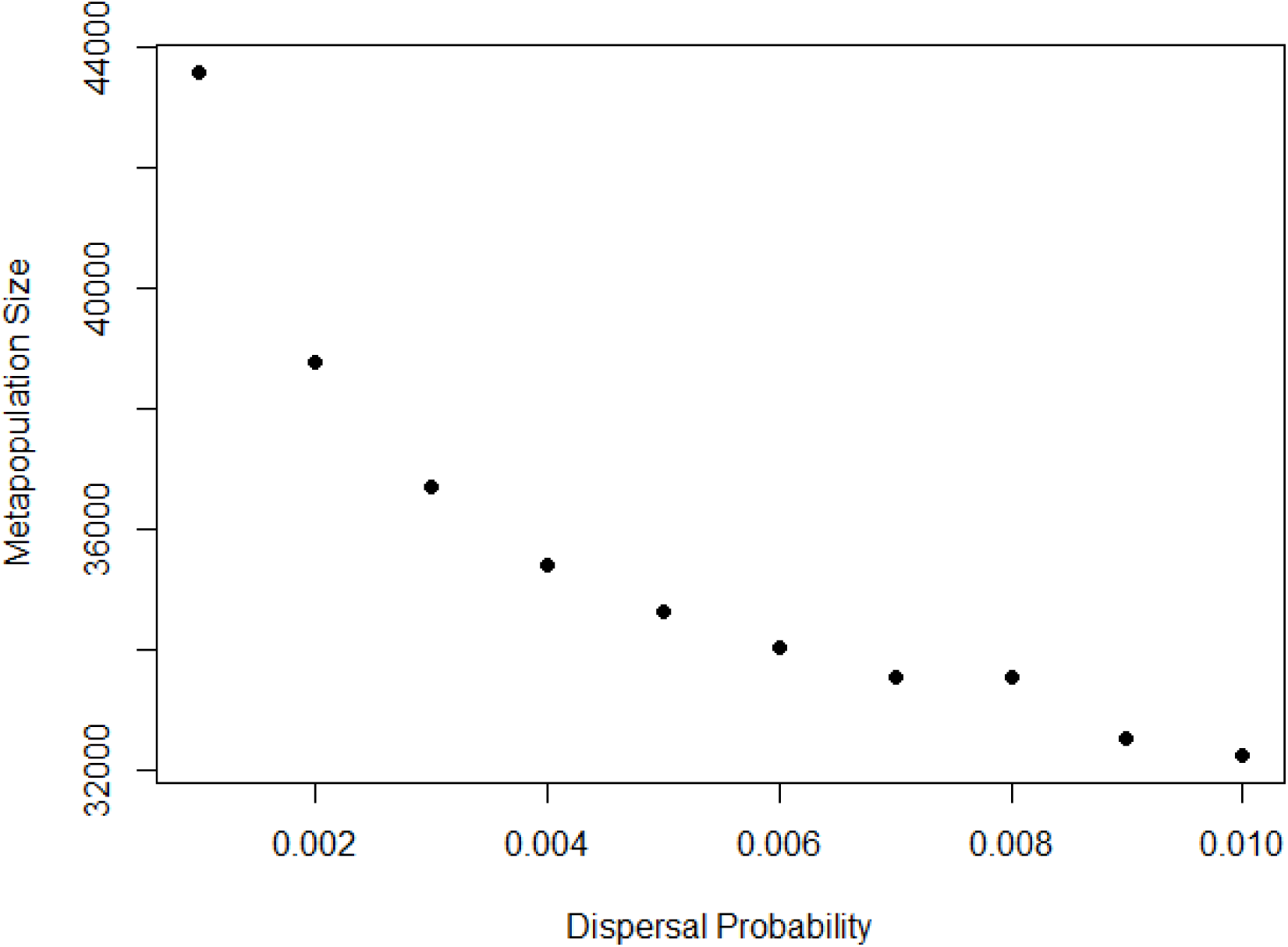
Metapopulation decline as dispersal probability increases in the absence of DFTD.

**Supplementary figure S4.**
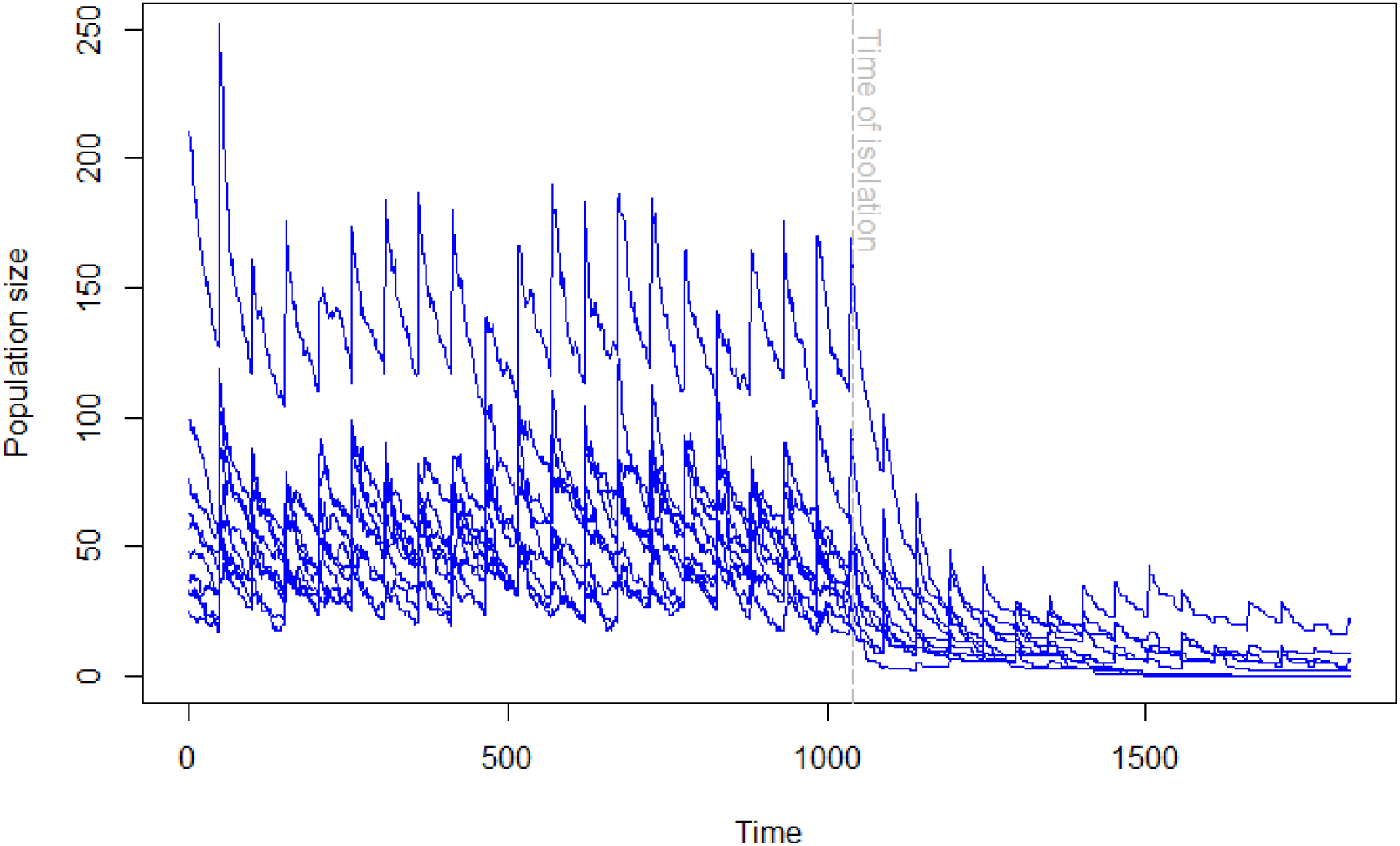
Example of the population decline observed in local populations after they are isolated from the rest of the metapopulation.

**Supplementary Figure S5:**
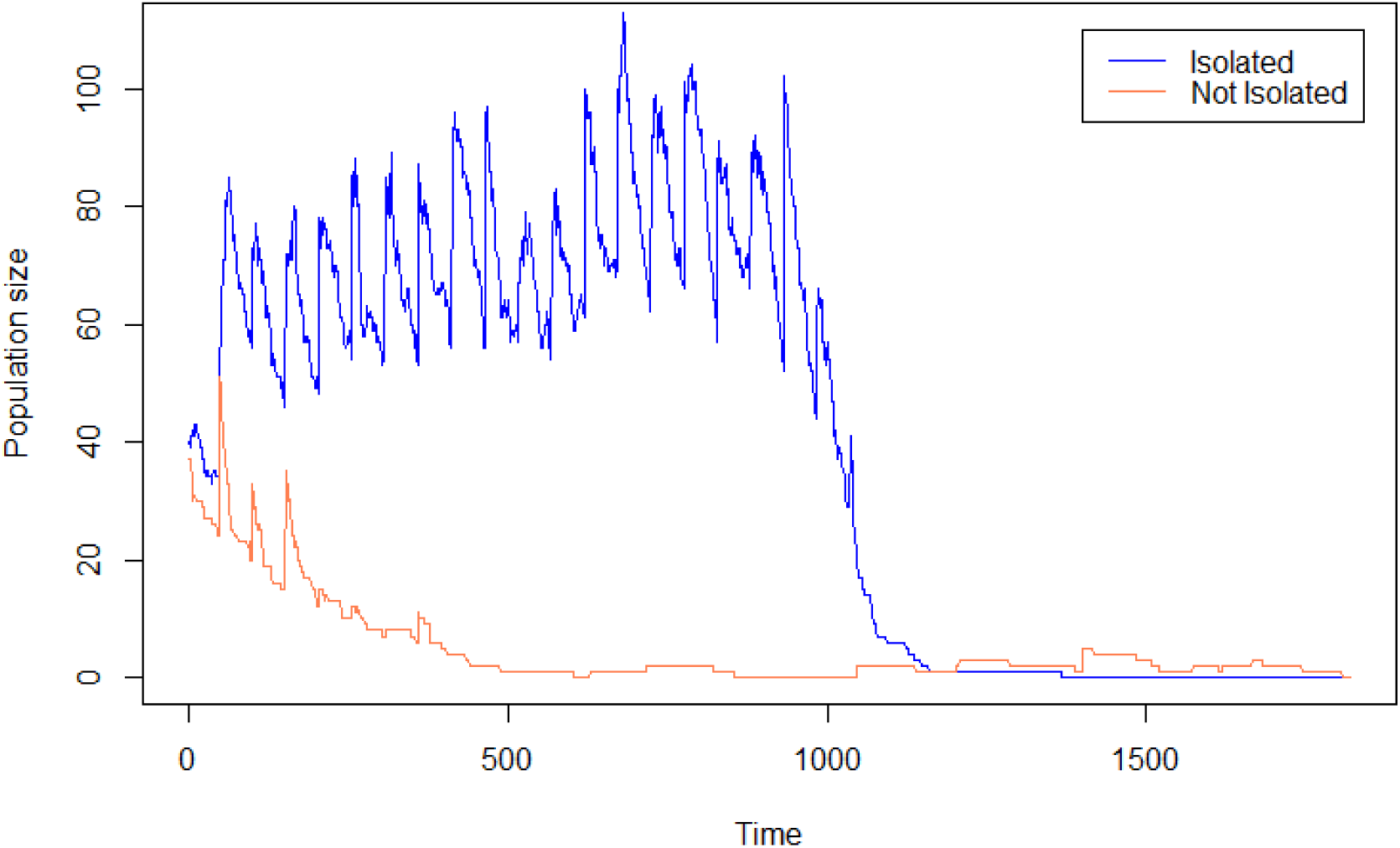
Comparison between isolated and non-isolated populations when the population size reduces to zero. The non-isolated population can become recolonised (but continues in an extinction-reintroduction cycle), whereas the isolated population cannot become repopulated.

**Supplementary Figure S6:**
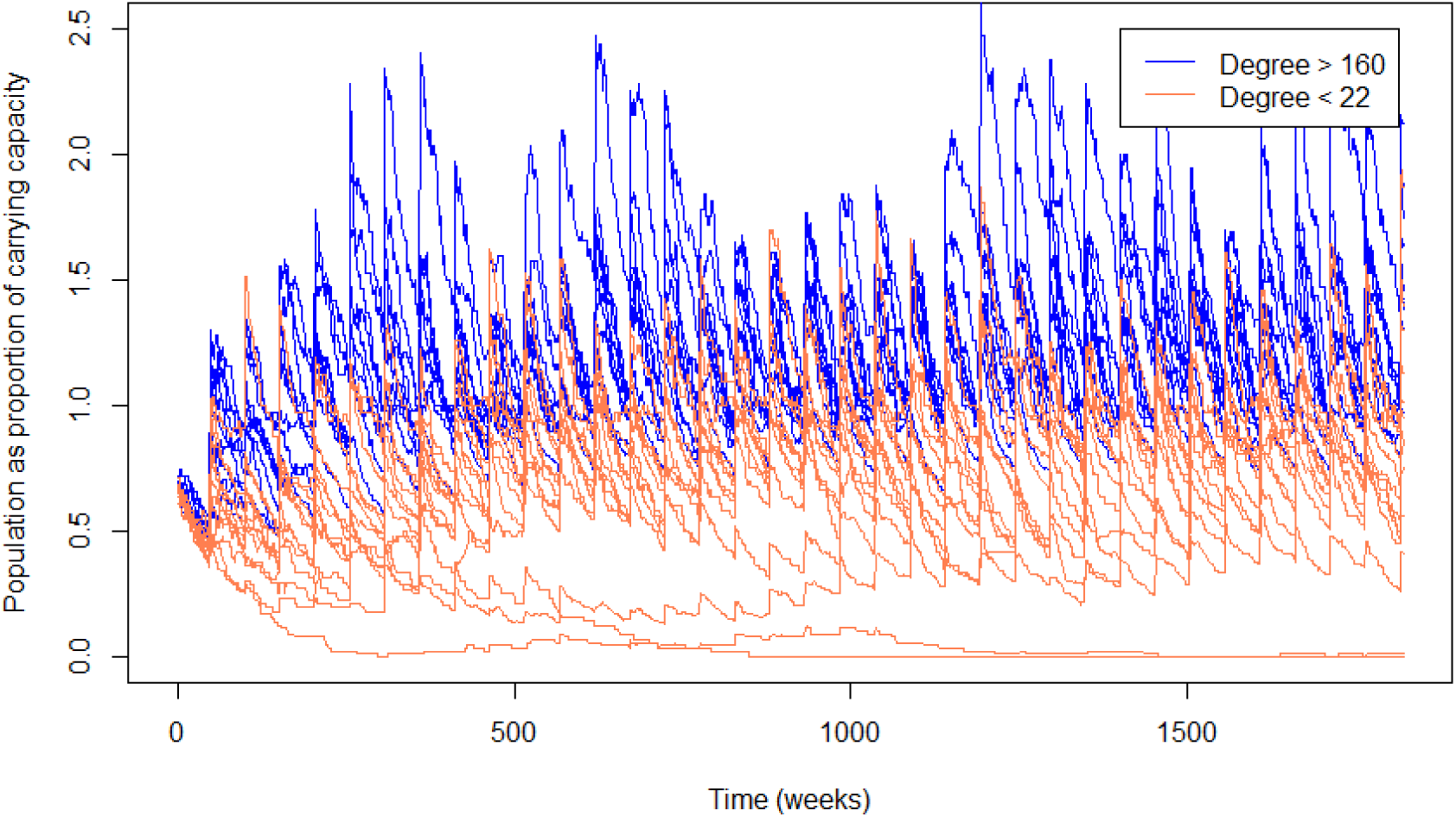
Example comparison of population size relative to carrying capacity for the most and least connected populations over time. More highly connected populations are consistently above their carrying capacity, suggesting that they are acting as pseudo-sinks, as suggested in (Zamberletti *et al*. 2018).

